# London taxi drivers exploit neighbourhood boundaries for hierarchical route planning

**DOI:** 10.1101/2024.02.20.581139

**Authors:** Eva-Maria Griesbauer, Pablo Fernandez Velasco, Antoine Coutrot, Jan M. Wiener, Jeremy G. Morley, Daniel McNamee, Ed Manley, Hugo J. Spiers

## Abstract

Humans show an impressive ability to plan over complex situations and environments. A classic approach to explaining such planning has been tree-search algorithms which search through alternative state sequences for the most efficient path through states. However, this approach fails when the number of states is large due to the time to compute all possible sequences. Hierarchical route planning has been proposed as an alternative, offering a computationally efficient mechanism in which the representation of the environment is segregated into clusters. Current evidence for hierarchical planning comes from experimentally created environments which have clearly defined boundaries and far fewer states than the real-world. To test for real-world hierarchical planning we exploited the capacity of London licensed taxi drivers to use their memory to construct a street by street plan across London, UK (>26,000 streets). The time to recall each successive street name was treated as the response time, with a rapid average of 1.8 seconds between each street. In support of hierarchical planning we find that the clustered structure of London’s regions impacts the response times, with minimal impact of the distance across the street network (as would be predicted by tree-search). We also find that changing direction during the plan (e.g. turning left or right) is associated with delayed response times. Thus, our results provide real-world evidence for how humans structure planning over a very large number of states, and give a measure of human expertise in planning.

## Introduction

Being able to plan for the future is a critical cognitive ability for humans. How humans might enact spatial planning has been the topic of a wide range of research (Hartley et al., 2003; Chrastil et al., 2015; Spiers and Gilbert, 2015; Weisberg & Newcombe, 2016; Horner et al., 2016; Epstein et al., 2017; Newcombe, 2018; Ekstrom et al. 2018; Brown & Chrastil, 2019; Miller & Venditto, 2020; Patai & Spiers, 2021; Matter and Lengyel, 2022). One common approach is to consider tree-search algorithms. Computationally, tree-search algorithms operate by repeatedly sampling different alternatives at each decision point according to some heuristic strategies and effectively compare entire sequences of decisions (Shannon, 1950; Daw, Niv, & Dayan, 2005; Huys et al., 2012; van Opheusden et al., 2021). More recent approaches involve Monte Carlo tree-searches that randomly sample trajectories potentially guided by state evaluation functions (Browne, et al. 2012), pruning strategies that eliminate unfavourable options (Huys et al., 2012), and reinforcement-based strategies that estimate the overall likelihood of a route being chosen (O’Doherty, Lee & McNamee, 2015; Botvinick, Niv & Barto, 2009). These models have managed to describe quite closely human and animal trajectories in small-scale and virtual reality environments (Daw et al., 2011; Russek et al., 2017; Miller & Venditto, 2020; de Cothi et al., 2022). Nevertheless, because the number of possible routes grows exponentially with each extra step to evaluate, tree-search becomes computationally intractable in large-scale environments such as most cities (Gershman, Horvitz, and Tenenbaum, 2015).

A promising development that could measure up to the computational demands of real-world navigation is hierarchical route planning (McNamee, Wolpert & Lengyel, 2016; O’Doherty, Lee & McNamee, 2015; Botvinick, Niv & Barto, 2009; Tomov et al., 2020; Bast et al., 2016). Hierarchical models segregate the environment and represent it through smaller, distinct areas that contain local information about specific places. The distinct areas are referred to as clusters, and the specific places as states. Instead of involving the entire environment, computations are then restricted to a subset of clusters and states. Route planning is first carried out across clusters / subregions to select the relevant ones. Within each cluster, a particular route is planned using the limited number of states available. Ultimately, this results in multiple sequences of shorter routes. This makes intuitive sense when we consider that cities will lend themselves to clustering in the different regions that evolve within its layout (Lynch, 1964; Filomena et al., 2019; Farzanfar et al., 2022).

Empirical studies of human navigation in small regionalized virtual environments have provided evidence of hierarchical route planning (Wiener, Schnee & Mallot, 2004; Wiener & Mallot, 2003; Balageur et al, 2016). Experimental work has also shown that humans decompose tasks in optimal ways for planning action sequences (Solway et al., 2014). Evidence for hierarchical route planning has also come from examination of reaction times of participants planning routes in a fictitious transport network, where the number of trainline interconnections, rather than the number of stations, has a significant impact on planning times (Balaguer et al., 2016).

Real-world evidence for hierarchically structured route planning, however, is scant. Taxi drivers in Paris have been studied to explore how they represent the city and plan routes (Pailhous, 1969; Chase, 1983). While this has shown evidence for use of the major street network to apply to planning routes (Pailhous, 1969) and greater overestimation of distances when two reference places were separated by neighbourhood boundaries (Chase, 1983), the studies do not provide an explicit test of hierarchical planning, such as has been used in lab studies (e.g. Balaguer et al., 2016).

One possible approach to exploring hierarchical planning is in the mental simulations of real-world routes, which have shown evidence of clustering of locations into regions and compressed representations of walking times or distances (Bonasia, Blommesteyn & Moscovitch, 2015; Arnold, Iaria & Ekstrom, 2016; Jafarpour & Spiers, 2017; Brunec et al., 2017). Likewise, the duration of mental planning of real-world routes can indicate regional hierarchies in a spatial environment (McNamee, Wolpert & Lengyel, 2016). Nonetheless, these findings concern planning times for a route as a whole. What has not been obtained to date is evidence from step-by-step planning in real-world, ecologically valid settings. The importance of this lacuna comes to the fore when we contrast the small state-spaces employed in controlled experimental setups (e.g. Wiener, Schnee & Mallot, 2004; Wiener & Mallot, 2003; Balaguer et al, 2016; Muhle-Karbe et al., 2023) and large street networks of the real-world (Griesbauer et al., 2022a). Lab experiments have used bounded spaces with artificially imposed accentuated hierarchies. Real-world cities, on the other hand, lack any straightforward segregation into clusters, often with no straightforward boundaries (Greisbauer et al., 2022a). For instance, the subway network in the study by Balaguer and colleagues (2016) was designed so as to be hierarchically segregated, and human cognition apparently leverages this structure for planning. It remains unclear where similar response patterns would be evident for route planning in real large-scale urban environments that lack such sharp hierarchical segregation.

The lack of defined boundaries is one of three methodological challenges presented when testing for real-world evidence of hierarchical route planning. The other two are the participants’ ability to give precise route descriptions, and familiarity. One can reliably expect participants to remember stops in a relatively small subway network and give clear descriptions of possible routes, but for an urban street network spanning tens of thousands of streets, there are very likely significant gaps in the spatial knowledge of typical residents. Finally, participants will have varying levels of familiarity with different regions of existing urban settings. Familiarity has been shown to affect performance in spatial tasks (O’Neill, 1992) and to distort representations of distances within an environment (Jafarpour & Spiers, 2017).

Here, we surmount these challenges by testing for hierarchical route planning in London licensed taxi drivers. Licensed London taxi drivers, also known as London cabbies (hereafter referred to as London taxi drivers), are famous for their thorough training on “The Knowledge of London”, which covers both main and secondary roads within the six-mile area around Charing Cross in central London, UK (see Griesbauer et al., 2022b). Years of driving experience in London, accumulated during training, as well as post-qualification during work, make such taxi drivers excellent navigators of London, and in contrast to the general population, they are trained to recall routes in all areas of London by giving precise travelling instructions and specific street names for each step of a journey through London (Griesbauer et al., 2022b). In addition, taxi drivers have a familiarity with all areas of London. Previous research with London taxi drivers has provided insights into the dynamics of cognition, neural plasticity and brain function during driving and navigating London (Maguire et al., 2000; Maguire, Woollett & Spiers, 2006; Maguire, Nannery & Spiers, 2006; Spiers & Maguire, 2006a,b, 2007a,b,c 2008; see for review Griesbauer et al., 2022b). Here we probe their capacity for planning. Towards this we have identified which boundaries in London are consistently recognised by London taxi drivers, e.g. Soho, Mayfair (Griesbauer et al., 2022a). Notably, many boundaries noted on official maps (eg. ‘The City of London’) were not consistently considered as boundaries. As a result, we were able to reliably segment London into different clusters that would be meaningful to this population.

We examined the response time to state the name of each street in the sequential planning of routes between a given origin and a destination pair (e.g. Kings Cross Station, to Paddington Station). If London taxi drivers employ hierarchical structures when planning routes, then response times should differ depending on the clusters they are planning over (streets on the boundary of regions vs other streets). If a tree-search is used to plan, the planning response time should be higher when there are more streets ahead to be evaluated as part of the plan. If they are exploiting boundary streets to plan they may be faster to respond when they need to select these streets as they would have been pre-selected in a hierarchical plan (McNamee, Wolpert & Lengyel, 2016).

The street network also contains additional features that may impact route planning. These include turning left or right, road classifications into main and minor roads, and Euclidean or path distance to the destination. Turns have been found to act as conceptual boundaries that have an impact on memory recall of route features (Lloyd, 1989; Kuipers, Tecuci & Stankiewicz, 2003; Brunec et al., 2020), on the estimation of path distances (Hutchenson & Wedell, 2009; Sadalla & Staplin, 1980; Brunec et al., 2017), and on preferences towards routes with fewer turns (Venigalla, Zhou & Zhu, 2017; Broach, Dill & Gliebe, 2012; Elliot & Lestk, 1982). Thus, turns may likely lead to slower response times. As for major roads (e.g. UK ‘A roads’), they might be better remembered than minor roads due to increased familiarity, as was the case with Parisian taxi drivers (Pailhous, 1969), which would favour a faster recall. Alternatively, major roads are associated with higher betweenness centrality in the street network (Javadi et al., 2017) and thus more long-range potential options for a plan, leading to predicted slower response times. Past research indicates distortions in the representations of space when a path must circumnavigate a region leading to high circuity metrics in a path (e.g. long path distance to a destination, but a short Euclidean distance) (Brunec et al., 2017). Thus, it seems possible that response times may be impacted by the degree of circuity of routes (e.g. straight routes vs U-shaped routes).

## Methods

### Participants

44 licensed London taxi drivers were tested after written informed consent. One participant displayed extremely unstructured route recalls that did not allow for a transcription of the routes, thus data from 43 taxi drivers (41 males, 2 females) was analysed. Their mean age was 53.82 years (*SD* = 10.35; range: 34-75 years) and their mean experience driving a taxi was 19.61 years (SD = 15.69). 19 taxi drivers participated in the first round of data collection (all male, age = 52.94, SD = 9.81, experience = 19.97, SD = 16.52) and 24 taxi drivers at the second round of data collection (22 male, 2 female; age = 54.50, SD = 10.93; experience = 19.30, SD = 15.32).

All of the taxi drivers were native English speakers. Data was collected in two periods with an interval of six months between the collection times. All procedures were approved by the ethics committee (CPB/2013/150 and EP/2018/008).

### Materials

The task for the first data collection consisted of 12 origin-destination pairs (e.g. from Chelsea harbour to London heliport). For the second data collection, two origin-destination pairs of the original set remained and 6 new pairs were added which increased the variety of route planning problems across London. From the two routes that remained the same, one origin location had to be updated (from Joe Allen’s restaurant to Bill’s) because the restaurant had moved location. The new origin (Bill’s) was a neighbouring restaurant to the original (Joe Allen’s restaurant). All pairs were chosen to vary in their geographical properties (e.g. path length, Euclidean distance, direction of travel; see Table S1). Euclidean distance was designed to be relatively decorrelated from the number of streets to the goal and potential boundaries that had to be crossed. This was to allow us to determine whether each type of distance impacted planning. For instance, route 7 and 18 had a similar number of streets (8 expected streets for route 7 and 7 streets for route 18), but varied in their planning distance (∼11km for route 7 and ∼1.5km for route 18). On the other hand, route 7 and route 17 were similar in their planning distance (∼11km vs ∼8.4km) but varied in the number of streets that had to be recalled (8 streets vs 14 streets). Similarly, the number of boundaries that we expected to affect routes varied across routes: Some routes did not cross any boundaries (e.g. route 4, 6 or 10), while other routes required at least partially to mentally travel along boundaries (e.g. route 7, 12 or 14) or crossing several boundaries (e.g. routes 1, 5, 13, 16 or 17). In collaboration with a London taxi driver training school (See Greisbaur et al., 2022b for details), teachers provided feedback to ensure the validity of the selected route with regards to route planning properties. Two SONY ICD-PX240 Mono Digital Voice Recorders were used in this experiment. One of the recorders was used to replay pre-recorded instructions and the route planning tasks. With the second recorder, the experimenters recorded the whole duration of the experiment, from the initial task presentation to the final route recall.

### Experimental Design

Licensed London taxi drivers were recruited in the area of Bloomsbury and the borough of Camden, London (UK). Before participating in the study, taxi drivers gave written consent and filled in a personal questionnaire to indicate age, gender, experience, whether they were native speakers, and whether they had taken part in this study on a previous occasion. After participating, taxi drivers received monetary compensation.

The group of taxi drivers who participated in the first study were verbally informed that they were to plan routes (*runs*) between origin-destination pairs (*points*) and that these routes would be presented through audio recordings. Participants were asked to listen carefully to the instructions, and they were warned that repeating instructions was not possible. If they did not understand or know either of the points, they could skip the route or carry it out where they perceived the points to be. In order to avoid interfering effects from obstructions unrelated to street network properties, participants were instructed to disregard possible congestion and any temporary obstructions in the street network. As some traffic rules can change in dependence of day and time, participants were prompted to imagine they were carrying out the route planning task on a typical Monday morning around 11.00 AM, to keep conditions consistent. Finally, in order to avoid distractions from the planning process and to elicit a structured recall across all drivers, participants were instructed to focus on the route planning as if they were in a ‘knowledge examination’ situation (Griesbauer et al. 2022b) and to refrain from questions, comments or explanations.

Instructions to follow the structured recall as in the examination situation were given only before the first and the second route plan, but drivers received a reminder of day and time (i.e. Monday morning at 11.00 AM) before each route. These reminders had been audio recorded and were presented together with the audio recordings of the twelve route planning in the following format:

> “Please remember to do this under appearance conditions. So, no questions or clarifications, when you hear the points. If you’re unsure, start or go where you think it is, or skip the run.” (Before route 1 and 2)

> “Please remember, you’re doing the next run on a Monday morning around 11 am.” (Before each route)

> “Please call out the run from [Point, Street] to [Point, Street].” (Task presentation) Drivers listened to the set of instructions and then planned the route before moving on to the next set of instructions and route planning task (for an illustrative example of a recall block, see Fig. 1). The whole sequence from the first instruction to the final route recall was recorded on a second audio-recording device.

**Figure 1.**
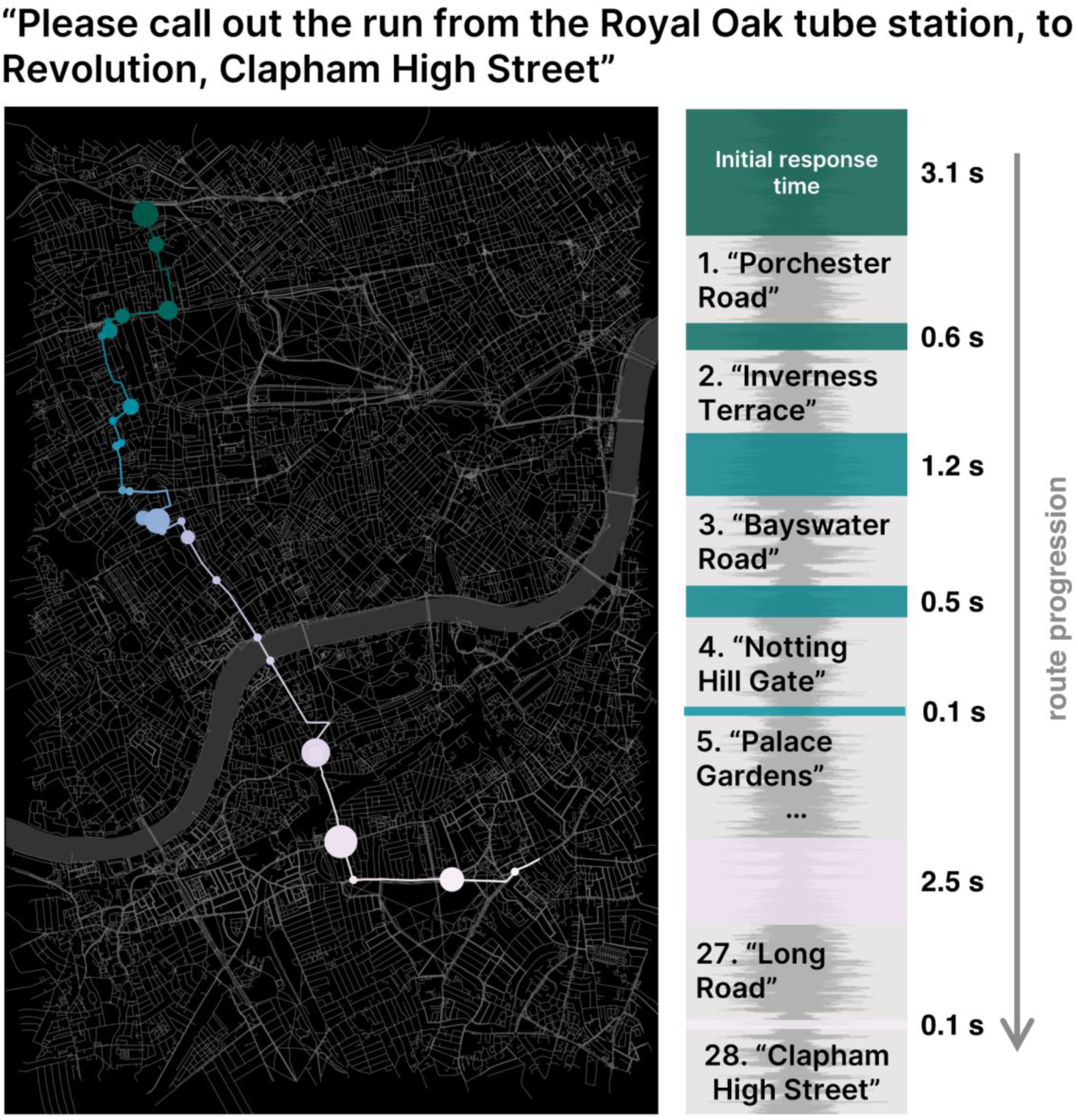
Example of one taxi driver’s plan for one of the eighteen origin-destination pairs probed. Taxi drivers received pre-recorded audio instructions of the origin and destination location and planned the route before being presented with the next route. All instructions and route recalls were audio recorded (top) and transcribed to extract response times and visualise planned routes (bottom right). The time between given instructions and the first recalled street was defined as the *initial response time*. Time between recalled streets of the route were defined as *response times between streets*. See Figure 2 for all the routes taken for this origin-destination pair. This figure was generated using Open Street Map data and the osmdata package in R.

For the second group of taxi drivers, who were presented with a modified set of eight origin-destination pairs, the following additional modifications were put in place to improve the procedure: During the first period of data collection, several taxi drivers repeatedly asked for clarifications of the origins or destinations or could not remember which location was named. Thus, several routes were skipped by drivers. Therefore, ‘flashcards’ were provided stating the location and the corresponding street for both origin and destination. These flashcards were shown to the drivers directly after they had listened to the audio recorded task and stayed visible during the entire route recall of a task. Additionally, after completing the first and second routes, drivers were asked for confirmation that the recalled route reflected what they would have done on a Monday morning at 11.00 am, which all taxi drivers confirmed.

### Data Transcription

The collected audio data of the recorded route recalls was transcribed in terms of street names and response times. Initial response times (i.e. the time between task instructions and the first street named) reflected route planning behaviour that included a variety of actions and processes inconsistent across drivers or routes (e.g. affirmative questions concerning locations). These actions were not consistent across or within taxi drivers or routes, and could not be separated from each other analytically, because most planning was carried out silently to ensure natural planning behaviour of the drivers. Therefore, initial planning times have been excluded from the analysis of response times. In contrast, *response times between streets* (i.e. pauses between two consecutively named streets) were part of the sequential recall of street names and reflected planning behaviour directly related to each point of the recall process, not leaving opportunities for unrelated planning actions (see Fig. 1 & 2).

The transcription of all response times was carried out with the free and open-source audio software Audacity, versions 2.2.2 and 2.3.1, which allowed for an accuracy of up to 0.1s. The street names were transcribed and corrected for mistakes (e.g. “Townsend Rd” to “Townmead Rd”) or unified (e.g. “Charles the 1st” to “Trafalgar Square”) to ensure comparability of response times at each street or place. The analysis of the complete dataset was carried out in R (version 4.0.2). To ensure the reliability of transcribed response times, an inter-rater reliability test was carried out for both studies. The intraclass correlation coefficient (ICC) for the first study (transcribed by two coders), was assessed through a two-way, mixed effects, absolute agreement, single-measures model. The ICC (ICC = 0.98, p < .001) was in the excellent range, suggesting a similar transcription of response times due to a high agreement between the two coders. For the second study (transcribed by four coders), a one-way random effects model with absolute agreement was used. The ICCs indicated a moderate range of agreement (ICC = 0.61, p < .001), which was considered acceptable for a rating involving four coders. A Wilcoxon Signed-rank test indicated no group differences between the two sets of taxi drivers for routes 7 and 8 for log-transformed (Mdn(S1) = −0.22, IQR = 1.18, Mdn(S2) = −0.16, IQR = 1.24, *p* = .252, r = .046) or z-standardised (Mdn(S1) = −0.29, IQR = 1.11, Mdn(S2) = −0.33, IQR = 1.24, *p* = .688, r = .016) response times between streets. Accordingly, the data from both instances of data collection was treated as one data set.

### Data Analysis

We used a linear mixed effects model to test the effect of street network variables on the log-transformed response times (c.f. Coutrot et al., 2018). Taxi drivers and their routes were entered as random effects to account for individual differences and potential correlations between repeated measures. The fixed effects variables of the model were *boundaries (B)*, *turn actions (T)*, *number of streets (N)*, *road type (R)* and *Euclidean distance (E)*. Here, the *boundaries* reflected agreement rates across taxi drivers in percentages (based on Griesbauer et al., 2022a). The *number of turns* was extracted from the route recall. Turns were coded categorically as *turn* where a change in direction occurred between consecutive streets or as a *forward* action where streets continued straight without a change of direction. The *number of streets* to the destination was counted down, with the last street having the value *1* and the first street the value *n*, if a driver recalled *n* numbers of streets for a route. *Road type* was categorised as either a main road (either trunk roads or primary roads), or other roads (i.e. all remaining road types, such as secondary, tertiary, or residential roads). Here, data about road type classifications was extracted from the OS MasterMap Integrated Transport Network (ITN) Layer (2018). *Euclidean distance* to the destination was calculated from each intersection of two consecutive streets to the destination. Data was extracted from OS MasterMap Integrated Transport Network (ITN) Layer (2018). These five fixed effects were decorrelated as all variance inflation factors were below 2.5. The basic model, which was used to describe log-transformed response times between streets, had the following structure:

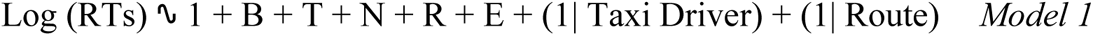

*Model 1* was designed to provide an initial assessment of the data based on factors found in the literature. Follow up analysis was carried out to examine in more detail the effect of boundaries, and two alternative models were considered. The first model (*Model 2*) replaced boundary streets with all the streets in the London district of *Soho* (*S*), including its boundaries. We chose to focus on Soho because it was the only district that conceptually appears as an ‘island’ with sharply defined boundaries in the central London street network (Griesbauer et al., 2022a) and appears to be treated distinctly by London taxi drivers (Maguire, Nannery & Spiers, 2006). In contrast, Mayfair and Belgravia are not entirely surrounded by other urban areas because they share boundaries with Hyde Park, a green space that is conceptually different from urban spaces. The second alternative model (*Model 3*) examined the impact of *circuity*, a similar concept to U-turn costs. This analysis was carried out because Balaguer and colleagues (2016) had found a cost of U-turns where participants had to head back along a subway line to reach the destination. Here, circuity was defined as the fraction of path distance to the destination divided by Euclidean distance to the destination, both calculated from each street. The closer the circuity value to 1, the more similar is the travelled route (path distance) to a straight line (Euclidean distance). The larger the circuity value, the more deviation there is of the path from a straight line. Since circuity and Euclidean distance to the goal were not independent from each other, Euclidean distance was replaced by circuity in Model 3. Hence, the two alternative models were (changed variable in bold):

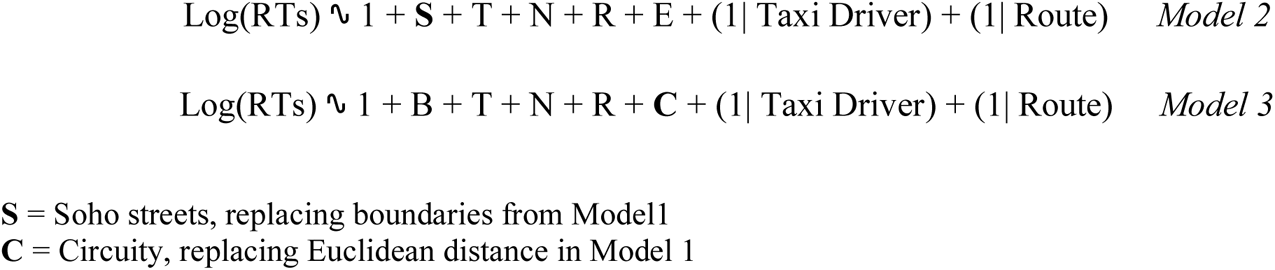

## Results

### Overview of the Data

The data, which was collected from *N* = 43 licensed London taxi drivers (green badge holders), included the recall of 354 routes (first study: 173, second study: 181). The mean for *initial planning times* was M = 13.83s (*SD* = 13.40; Table 1) over N = 315 routes. Data of some initial planning events had to be removed as drivers asked for clarifications and tried to engage in conversations. On average, taxi drivers recalled 9.1 out of 12 routes during the first study, and 7.5 out of 8 routes during the second study.

**Table 1.** Variable Summary for recalled data.

|  | **N<sub>1</sub>** | **Mean** | **SD** | **Min** | **Max** |
| --- | --- | --- | --- | --- | --- |
| **Initial Planning Time<sup>2</sup> (sec)** | 315 | 13.83 | 13.40 | 1.00 | 102.00 |
| **Mean Response Times between Streets<sup>2</sup> (sec)** | 3398 | 1.82 | 3.24 | 0.10 | 61.4 |
| **Total Response Duration<sup>2,3</sup> (sec)** | 354 | 17.53 | 16.47 | 0.20 | 89.40 |
| **Total no of Streets Recalled** | 354 | 10.71 | 5.71 | 3.00 | 30.00 |

<sup>1</sup> Number of occurrences in data set
<sup>2</sup> Minimum coding accuracy of response times: 0.1s
<sup>3</sup> Mean of the sum across all response times between Streets per taxi driver and per planned route

There were a total of 3398 responses, with a mean *response time between streets* of M = 1.82s (*SD* = 3.24). The *total response duration* for routes, a measure to reflect the total planning duration based on responses between streets per driver and route, was M = 17.53s (*SD* = 16.47). See Figure 2 for an example of the routes chosen for an origin-destination pair. For the raw dataset, these response times between streets were skewed towards minimal response times (see Fig. S1). Log-transformation (using the natural logarithm) and z-standardisation provided a better fit with a normal distribution, with the exception of high density at minimal response times of the log-transformed data. This was also reflected across routes (see Fig. S2). Mean response times between streets ranged from 0.7s for route 10 to 2.8s for route 9 and violin plots indicated skewness of raw data towards fast recalls between streets. These were ordered by ascending means of z-transformed data to allow for comparison across routes whilst accounting for individual differences between drivers. A high number of outliers for the raw and z-transformed data at each route highlighted slow responses between named streets with up to 60s. Log-transformed violin plots showed two high density peaks of data, one near fast response times around 0.1 s (log-values around −1) and a second density peak near values of 1s (i.e. log-values around 0). For this study, the high density of fast response times (around 0.1s) was thus considered as a measurable lower bound on response times from the audio recordings. In contrast, deviations towards slower responses (i.e. second density peak of log-transformed data and outliers in the raw and z-transformed data set) were expected to convey information about spatial structures indicative of potential planning of new sequences (and therefore of hierarchical planning). Similar to response times between streets, the means of response times between streets by route, the total number of recalled streets per route, and initial response times were all left-skewed for raw data. After log-transformation and z-standardisation initial response times fit a normal distribution (see Fig. S1). No relation was found between raw or transformed initial response times and mean response times between streets. Thus, there was no evidence that those who engaged in longer initially thinking about the route were then faster at calling it out street-by-street.

**Figure 2.**
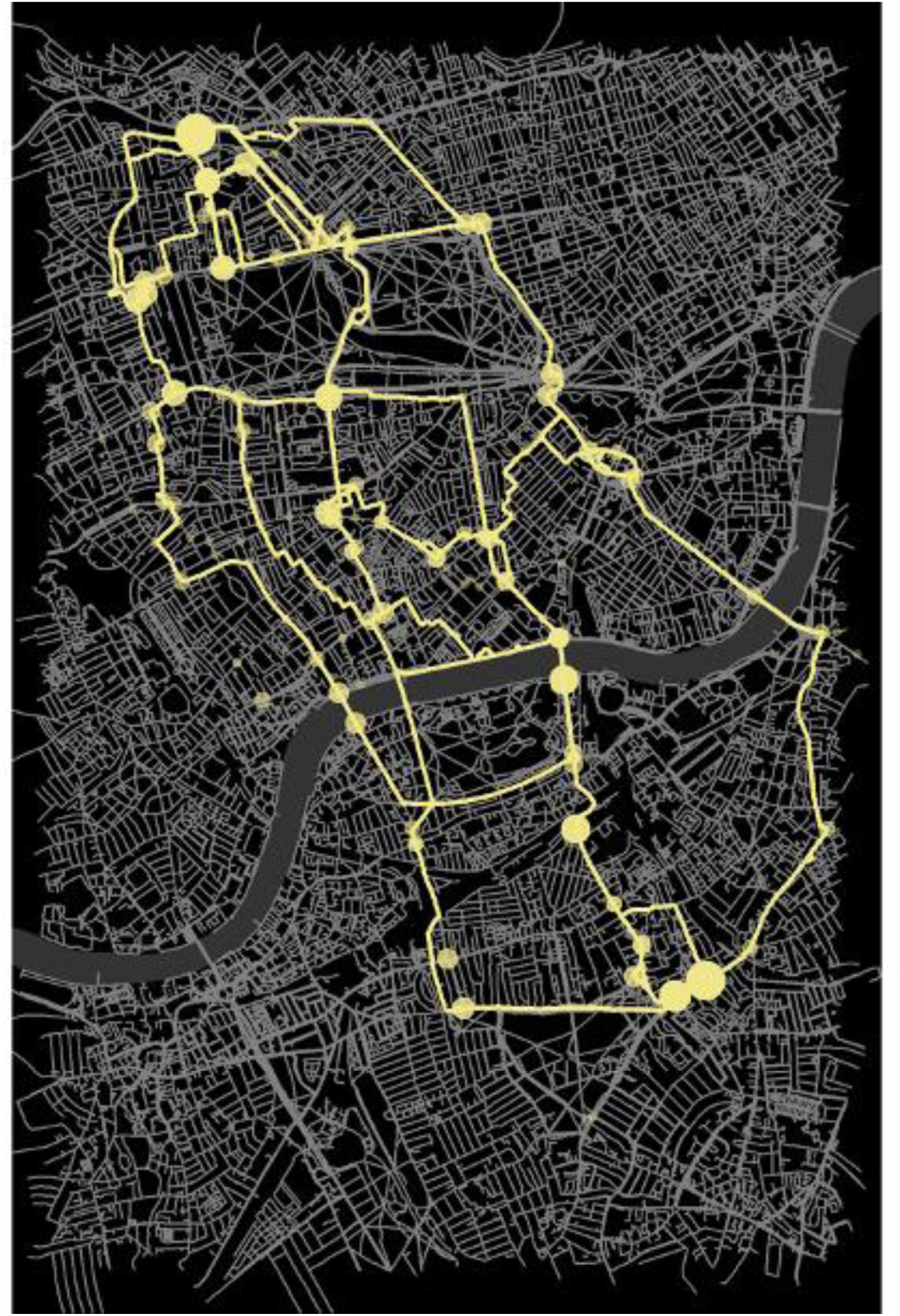
Overlay of the routes planned by all taxi drivers for a single origin-destination pair. Taxi drivers received pre-recorded audio instructions of the origin (Royal Oak tube station) and destination (Revolution, Clapham High Street) locations and planned the route. All instructions and route recalls were audio recorded and transcribed to extract response times and visualise planned routes (in yellow). Time between recalled streets of the route were defined as *response times between streets*. Larger circumferences on the map (in yellow) correspond to longer response times between streets. All the responses from all participants for a single origin-destination pair have been overlaid on the map of London. This figure was generated using Open Street Map data and the osmdata package in R.

### Age and Experience

We considered that with experience taxi drivers might improve with their navigation, and conversely with age they might decline (see e.g. Coutrot et al., 2018). Thus we studied the relationship between *age* (M = 53.82 years, *SD* = 10.35) and *experience* driving a taxi (M 19.61 years, SD = 15.69) for the group of 43 taxi drivers (see Fig. 3). A Spearman correlation indicated a strong, significant positive relation of age and experience (*r_s_ (43)* = .73, *p* < .001). No relation was found between means of log-transformed response times by route and experience (r_s_ (40) = .08, p = .623) or age (r (37) = .13, p = .445) for the entire group of taxi drivers. Age and experience were decorrelated in a group of taxi drivers with 25 years of experience or less (r (24) = .23, p = .564). In that sub-group of taxi drivers there was also no relation between means of log-transformed response times by route and experience or age (experience: r (25) = -.11, p = .584; age: r (24) = -.11, p = .603). Thus, when separating age and experience, we find no evidence of decline with age and no gain with experience.

**Figure 3.**
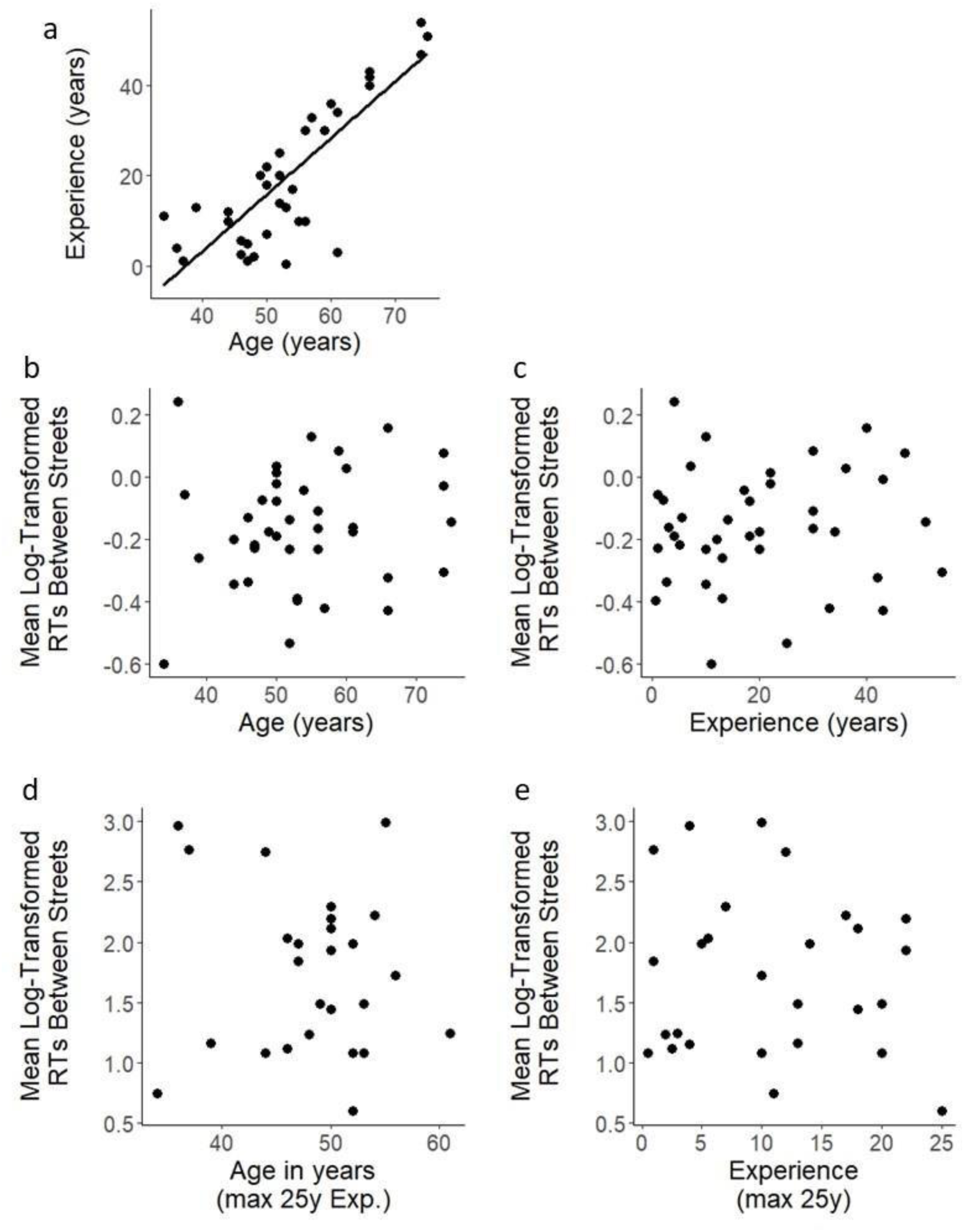
Age-experience relationship. Age and experience across taxi drivers was strongly correlated for the entire group of participants (a), but decorrelated for a subset of drivers with 25 years of experience or less. Neither the entire group of taxi drivers (b, c) nor the subset of drivers with 25 years of experience or less (d, e) showed any relation to mean log-transformed response times between streets.

### Turns, boundaries and number of streets to the goal impact response times

To see examples of scaled response times for routes where many taxi drivers chose the same path, see Figure 5. Examining the responses across these routes shows consistent patterns of speeded responses and slow responses. To examine the impact of different variables on the response times we analysed the data over all the routes, where sufficient variation was present across variables to estimate their unique contribution. Model 1 revealed that *boundaries* (b = - 0.082, *p*<.05, 95% CI= - 0.047- −0.12), *turns* (b = 0.13, *p* < .001, 95% CI= 0.17- 0.093), *number of streets* to the destination (b = 0.011, *p* < .05, 95% CI= 0.040 - −0.018), and *main roads* (b = 0.089, *p* <.05, 95% CI= 0.12 - 0.054) but not *Euclidean distance* (b = −0.038, *p* = .126, 95% CI= 0.013 - 0.063) had significant effects on *response times* (Fig 4). Coefficients of the model are reported in Fig. 4a, these parameters were additionally visualised individually in Fig. 4b-f. Thus, streets that required a turn into, or that were main roads, were associated with slower response times (Fig. 4). Streets classified as boundary streets had a faster call out than other streets (Fig. 4d). There was also a small but consistent effect that the greater the number of streets to the destination the slower response times (Fig. 4a,e). This difference in response speed was most evident for initial streets and final streets (Fig.4e), but not strongly evident when the full range of streets was examined (Fig. 4f). Euclidean distance did not contribute to the outcome of the model. Thus, it is not the distance ‘as the crow flies’ that will determine planning speed, but rather the number of streets to be travelled.

**Figure 4.**
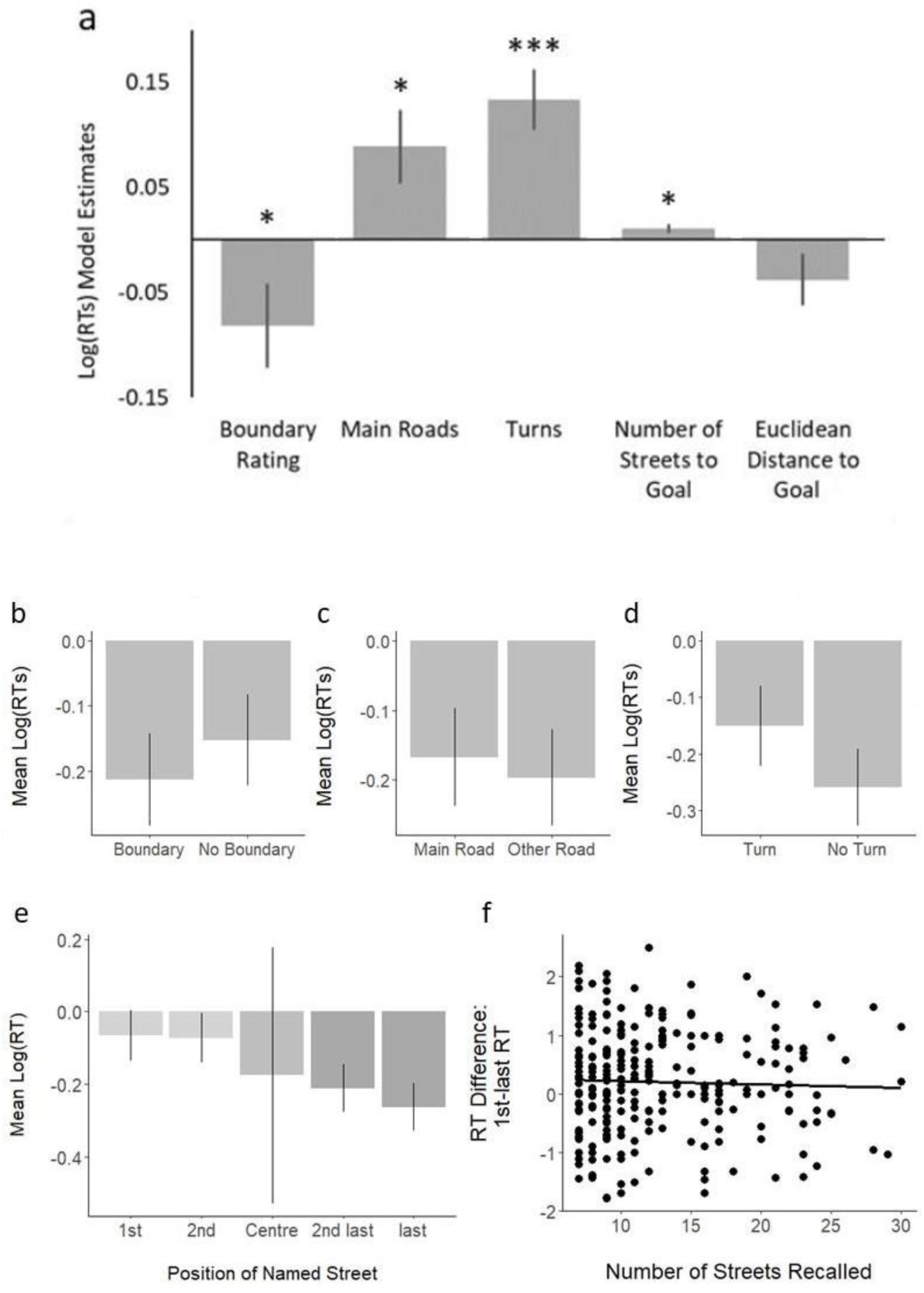
Evidence for hierarchical planning revealed by faster responses on boundaries. Linear mixed model estimates (a) indicate a highly significant effect (* < 0.05, *** < 0.001) of turns, main roads, the number of streets to the goal and boundaries significantly impact on log-transformed response times. Euclidean distance does not reach significance, nor did circuity when used in the model (model 3 see methods). (b-d) Differences in the recall of turns and non-turns (b), main roads and minor roads (c) and boundary streets and non-boundary streets (d) are plotted to allow visualisation of the response patterns. Additionally, the position of the street along the sequential recall highlights differences between initial and final streets (e). The difference between the first and last street is plotted for individual responses for routes containing 3 to 30 streets, revealing little evidence of slowing when the route planned is extensive (e.g. 30 streets) vs when it is short (e.g. 3 streets).

**Figure 5.**
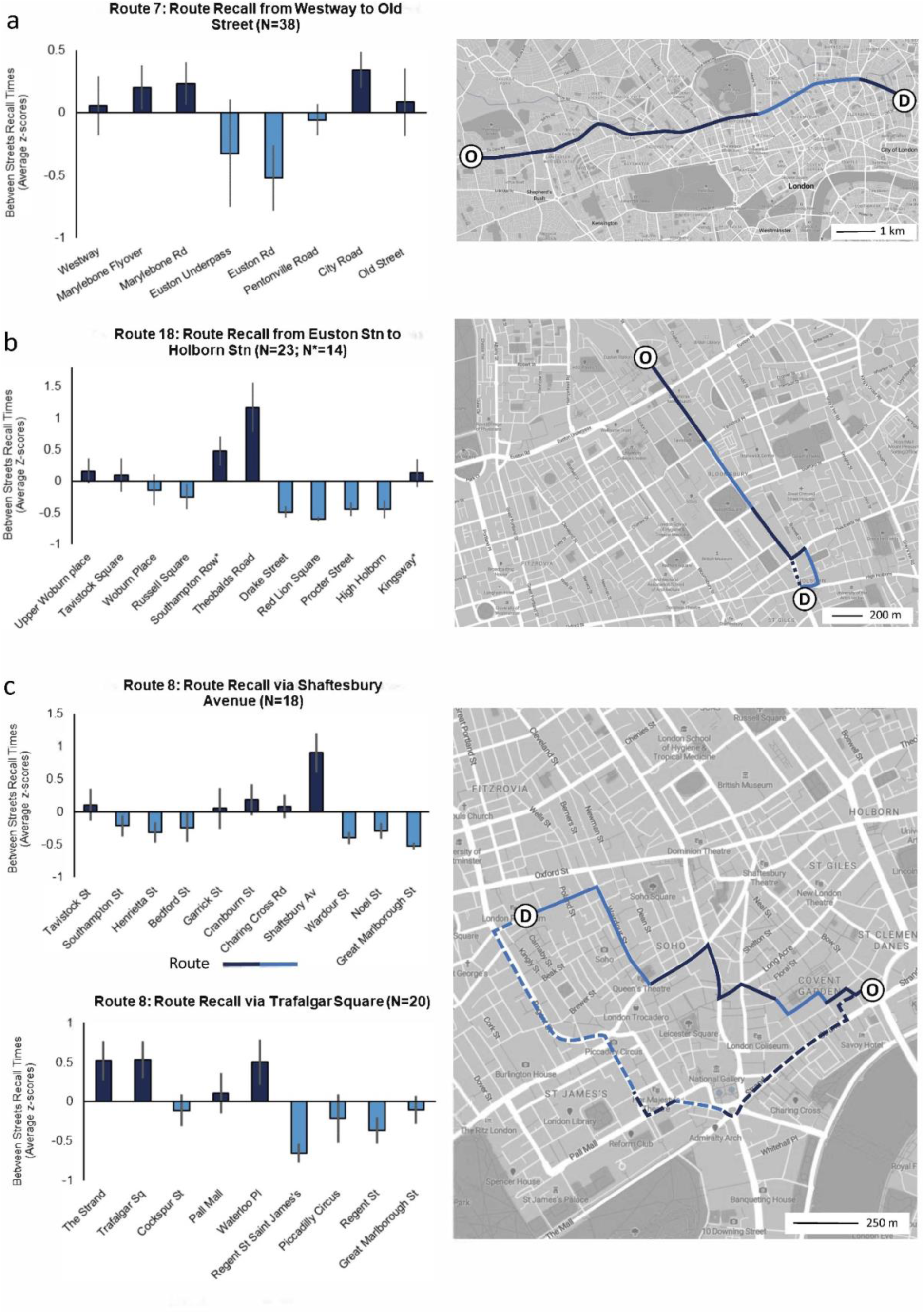
Routes where taxi drivers chose very similar routes. Taxi drivers planned routes between origin (O) and destination (D) pairs. A selection of three routes with a high agreement on the route across drivers is displayed to highlight differences in z-standardised response times between streets. Agreement was found for linear routes (a, b) and routes with two alternatives (c). Corresponding streets with faster (light blue) and slower (dark blue) than average recall speed are highlighted on the right. Note that in (b) only nine drivers planned to do a loop past Red Lion Square, the other 14 taxi drivers planned to go straight from Southampton Row to Kingsway. Map source: Mapbox.

Next we focused our analysis on the most consistent bounded region of London: Soho (Griesbauer et al., 2022a). Model 2 revealed significant effects of *Soho* streets (b = −0.16, *p*<.05, 95% CI= −0.094 - −0.23), *turns* (b = 0.14, *p* < .001, 95% CI= 0.17 - 0.11) and *number of streets* to the destination (b = 0.0095, *p* < .05, 95% CI= 0.014 - 0.0052). The effect of *main roads* was marginally significant (b = 0.089, *p* < .05, 95% CI= 0.95 - 0.029) and *Euclidean distance* was again not significant (b = −0.037, *p* = .135, 95% CI= −0.012 - −0.061). We found the response times of streets recalled within Soho were faster than for the rest of the environment. All other fixed effects variables indicated similar results, except or main roads, which only reach marginal significance. The second alternative, Model 3, examining *circularity* revealed that this variable was not significant (b = 0.014, *p* = .272, 95% CI= 0.026 - 0.001), see Fig. S3. In this model there was again a significant effect of *boundaries* (b = −0.085, *p*<.05, 95% CI= −0.045 - −0.12), *turns* (b = 0.14, *p* < .001, 95% CI= 0.17 - 0.11), *number of streets* to the destination (b = 0.0061, *p* < .05, 95% CI= 0.0091 - 0.0032), and *main roads* (b = 0.084, *p* < .05, 95% CI= 0.12 −0.049). Thus, from our extended models we show that planning across a set of streets in a well known bounded region is faster compared to other streets (matching predictions) and that routes that include higher circuity were not slower to provide responses in planning (not matching predictions).

### Streets with longer names did not have longer response times

We considered the possibility that it could take longer to recall the exact name of street names which are longer (e.g. ‘Stoke Newington Church Street’, is long, while ‘Oak Row’, which is much shorter). We found no relationship between the two normally distributed variables - short and long street names (see Fig. S5). A linear mixed model that additionally accounted for the number of characters as a fixed effect in the original model indicated no significant impact of the length of the street name (b = 0.0033, *p*=.410, 95% CI= −0.00071 - 0.0074). All other fixed effects variables were in line with previous model results (*boundaries*: b = −0.082, *p* < .05, 95% CI= −0.12 - −0.042; *turns*: b = 0.13, *p* < .001, 95% CI= 0.11 - 0.16; *number of streets*: b = 0.011, *p* < .05, 95% CI= 0.0062 - 0.015; *road type*: b = 0.087, *p* < .05, 95% CI= 0.052 - 0.12; *Euclidean distance*: b = −0.0384, *p* = .122, 95% CI= −0.063 - −0.014). Thus, the length of street names did not impact responses of the recall of individual streets.

## Discussion

Here we provide evidence of hierarchical route planning in a large-scale real-world environment. We asked licensed London taxi drivers to plan the route between a set of origin-destination pairs and recorded the time taken to state each street name (response time) they would use in the route. We tested the effect of spatially embedded street network variables on their response times using linear mixed effects models. Our results indicate a consistent pattern in modulations of response times in relation to spatial features, including the impact of hierarchical structures such as boundaries. We first discuss the impact of boundaries on response times and then consider the impacts of other variables, such as turns, circuity and number of streets in the planning process.

In contrast to prior studies of hierarchical route planning using artificial environments, which relied on visually distinct boundaries between districts / clusters in very small networks (e.g. Wiener & Mallot, 2009; Wiener, Schnee & Mallot, 2004; Schick et al., 2019), our study used boundary agreement rates for London districts to reflect the gradual differences in boundaries, characteristic of real-world cities (Cohen et al., 1978; Manley, 2014; Filomena et al., 2019; Griesbauer et al., 2022a). We found that taxi drivers were faster to select the next street when that street was a boundary rather than a non-boundary street. This finding appears to conflict with prior evidence from navigating fictitious subway networks, where responses are slower at stations on the boundary between subway lines (Balaguer et al., 2016). A plausible explanation is that, in subways, boundaries and the number of options for next states are confounded, whereas in real-world city streets, boundaries can occur across sets of streets with no change in the number of path options. We reveal that when it comes to navigating a large city, boundary streets are more quickly selected in the plan than other streets. This would appear to conflict with the idea that one might plan up to a boundary and then decide the next actions with a region; ‘fine-to-coarse’ planning with fine spatial information for the close environment and coarse spatial information for distant locations (Wiener et al., 2003; Spiers & Maguire, 2008). Rather our results are more consistent with taxi drivers having pre-planned a route before the step-by-step description given. In this plan, the boundary streets would be a prime consideration (McNamee et al. 2016). It is plausible that part of this hierarchical planning is explicitly considered by taxi drivers. This is because London taxi driver training schools use a method of training taxi drivers to consider sub-goals on their route when given a start and destination (Griesbauer et al., 2022b), and that such sub-goals correspond to boundary streets. However, future research will be required to explore the extent to which the initial planning period involved explicit consideration of boundary streets.

While the impact of boundaries provides support for hierarchical planning, our results do not negate a tree-search plan being deployed. Akin to Balaguer et al. (2016) we also found the number of states to the goal (in our case streets) was positively associated with response time. A tree-search approach to planning predicts slower responses (more options to consider) for states with more states to the goal (Streeter & Vitello, 1986; Elliott & Lesk, 1982; Miller & Venditto, 2020). Thus, our study provides both evidence for hierarchical and tree search planning. However, the impact of the number of streets to the goal was small. When we examined the response times as a function of the number of streets on the route (3 to 30) the impact was very minimal. Routes requiring responses with 30 states to reach the goal had similar response time differences between the first and last street as routes with 3 streets. This result strongly supports hierarchical planning as the dominant means by which London taxi drivers compress their representations of London to aid planning.

### Impact of road network structure and turning on planning demands

Our finding that turning left or right caused slowing in response times is consistent with results from previous studies exploring the impact of turns on spatial memory (Brunec et al., 2020). This slowing is consistent with taxi drivers visualising the route as they progress, because visualising a turn would plausibly be more demanding than continuing straight ahead. A previous study in which taxi drivers gave post-task verbal reports of their thought process indicated that many taxi drivers, years after initial training, visualise the different streets to the goal as part of the planning process (Spiers & Maguire, 2008). Our investigation of the training process revealed that visualising the route was one of the key methods used to train taxi drivers in their spatial memory and learning of London (Greisbaur et al., 2022b). Notably, the response time for left and right turns is not trivially due to the additional delay that would be incurred by having to add the words “left into..” or “right into..” before street names; all responses required a statement about the direction, e.g. “forward into..” if there was no turn.

It could be expected that if a street is a major street in the network (e.g. Oxford Street), it would be easier to remember and thus quicker to be included in the plan. Indeed, Parisian taxi drivers have been previously reported to prioritise such streets for planning (Pailhous, 1969) and an amnesic London taxi driver was able to navigate by such streets, despite losing memory for minor roads (Maguire, Nannery & Spiers, 2006). However, this is not what we found. Rather taxi drivers were slower on average when a major road was the next step in the planned route. A plausible reason why major roads were associated with slow responses in the current study comes from the fact that such streets are associated with more long-distance connections than minor roads, as formalised by a higher betweenness centrality (for London, see Javadi et al., 2017). As such this means the number of possible routes from such a street to other locations is substantially increased, driving a greater demand on planning.

The only spatial variables that had no significant impact on response times were Euclidean distance and circuity (i.e. path distance to the goal divided by Euclidean distance to the goal). For many routes, Euclidean and number of streets to the goal are correlated. Here we designed routes that decorrelated these, helping to reveal the specific effect of ‘streets to the goal’ from ‘Euclidean distance to the goal’. The absence of an effect of Euclidean distance suggests that, while taxi drivers might visualise turns into new streets, or have higher demands in rotating travel in a mental model, they do not consistently create mental images of views along the entire real route to the goal. Rather it would be consistent with a sequence of mentally ‘snapshots’ street to street point for planning. This again mirrors the training of taxi drivers in which they learn a visual view for each street and exploit this for planning (Greisbaur et al. 2022b). The absence of circuitry on planning demands is surprising and suggests that it is the event of turning in a new direction that is demanding, and that once mentally simulating a route in a given direction, there is no specific mental cost to continuing in that direction, even though it may be away from the goal.

### Limitations

There are several limitations to our study. Firstly, we recruited drivers in the areas of Bloomsbury and King’s Cross, and they showed preferences for working in Central London, which could have impacted their knowledge of areas with greater distance to central London (e.g. West London, south of the River Thames). Future studies could recruit taxi drivers at different taxi ranks across London to account for this possibility. Secondly, even though the study included several geo-spatial properties (e.g. distance measures, boundaries, road types, turns), there is a range of information that we did not account for in the current analysis. For instance, planning directions, angular deviations at each street, spatial analytics or perceptual input of building use were not included (even though angular deviation and circuity are related metrics). These variables went beyond the scope of the study, but they should be considered in the future.

Potentially confounding factors, such as linguistic factors, age, or experience, were not found to impact response times. We considered whether the length of street names may interfere with response times during the verbal recall, but no such effect was found. For the entire group of taxi drivers, age and experience were correlated, and as age increased, so did experience. Age would be expected to impair route planning (van der Ham & Claessen, 2020), and experience should have the opposite effect (Pailhous, 1969; Chase, 1983). Moreover, training of spatial navigation abilities is expected to cause changes in hippocampal volume, the neural centre for spatial navigation: in a series of neuroimaging studies, volume changes in the posterior hippocampal grey matter were correlated with experience, and with greater experience one might expect better performance (Maguire et al., 2000; Maguire, Woollett & Spiers, 2006). However, the current data does not seem to point in this direction, as there were no correlations between experience and response times. We considered that for the entire group of taxi drivers, age and experience might have cancelled out any response time related effects. Even for a subset of drivers, where age and experience were decorrelated, no impact of age or experience was found. It is possible that the thorough training and daily use of navigation skills among London taxi drivers might protect them from age-related decline in spatial skills (Lövdén et al., 2012). Ultimately, this might explain why neither the entire group, nor the subset of taxi drivers showed any ageing effects for response times of route recalls.

While we controlled for word length, other linguistic factors, such as word complexity or familiarity of street names could have acted as confounds. Or taxi drivers might be aware of a street but struggle to recall it by name. Additional validation of findings could be achieved through alternative approaches with designs that do not rely on verbal recall of street names. For instance, video recorded route drawings on maps could provide supporting evidence for the current results. However, such a design would draw less on mental representations because visual features of the map (differently highlighted routes) are likely to impact route planning, given the availability of external spatial information. It is also not the manner in which taxi drivers are formally tested on competency. Therefore, study designs might have to restrict visual information to a small area around the streets that are being highlighted to ensure planning to rely on mental planning (e.g. showing only an area smaller than a quarter mile on a map at any time). In turn, implementing such a study would require technical or technological approaches that could come with other drawbacks regarding motor actions and a preference for paper maps over digitised maps with taxi drivers. Alternatively, drivers might be prompted to highlight key points that they would pass through on a map, or to indicate if specific streets would be part of their route. Under time pressure, these prompts might highlight response time differences for street network structures that are part of a hierarchical representation when contrasted with streets that are non-hierarchical. Here, planning would have to depend on predefined conditions, such as the most direct route rather than general preferences as in this study. However, such approaches would not allow for rich information on street level, focusing instead on coarse route level information, because attention is drawn to key points rather than entire routes. While each method comes with its own drawbacks, a diversity of approaches in future work might provide a useful contrast to the current study.

### Future directions

In this study, we tested a total of 43 taxi drivers across two sets of data collection, and we analysed a total of almost 3,400 response times between streets in relation to different street network properties. This data allowed us to identify interesting effects in the route planning process of licensed London taxi drivers, such as the impact of boundaries, turns or road type. There is room, however, for further spatial analysis of the environment under study. Space syntax, for instance, could provide vital insight into the role of major roads and street network properties in route planning (Peponis et al., 1990; Hölscher et al., 2006; Javadi et al., 2017; Yesiltepe et al., 2023). The current approach did not allow for this because most spatial network analytics use an approach that segregates entire streets into several clusters and then attributes spatial measures to each cluster, rather than to an entire street and their corresponding response times. Additionally, there is currently no approach to automatically assigning cluster-based information from the space syntax dataset of London to the street network, as clusters were parts of an unlabelled graph (i.e. no assignment of street names) and only contained graph network related information (e.g. betweenness centrality). Future analysis could consider a manual approach and focus on just a subset of routes or a specific area such as Soho.

Another question that remains open is how the spatial variables under study can be used to predict route planning at places in which multiple features interfere with each other. For instance, we would have predicted a fast recall for Shaftesbury Avenue in route 8, but the response time turned out to be extremely high. It appears that the properties that it was both a main road and a turning outweighed boundary factors in this particular case. Additionally, the intersection of Charing Cross Road with Shaftesbury Avenue might have turned it into a decision point, as both roads are main roads and several alternative options have to be considered, such as whether to cut through the north-eastern corner of Leicester Square. Thus, when forming predictions for a complex, real-world environment it will be useful to develop approaches to weigh many factors that might impact planning at specific places e.g. using judiciously chosen interaction variables for linear modelling, and this is an area that requires further research.

London taxi drivers are expert planners trained to navigate a particular urban environment. It is increasingly clear that there is a wide diversity of expert navigation strategies across the world (Fernandez Velasco & Spiers, 2023). It would be interesting to employ the methods developed here to investigate the factors that impact planning response times in other expert navigators (e.g. those navigating river system networks). Future work could also investigate more explicitly how navigation choices in the network related to metrics from reinforcement learning agents (de Cothi et al., 2022) or metrics about the space such as centrality measures (Lancia et al., 2023) and street network entropy (Coutrot et al., 2022).

Recently, there have been some important advances concerning the brain regions involved in hierarchical representations (Peer & Epstein, 2022; Andermane, Joensen & Horner, 2022). Expert planning of the kind studied in this paper is likely to involve prefrontal-hippocampal interactions (Simons & Spiers, 2003; Patai & Spiers, 2021). Previous work has shown differences in hippocampal engagement in familiar environments when many vs few turns are required to reach a goal (Patai et al., 2019). Lateral prefrontal activity has been found to scale with the number of future options in streets when re-planning was required (Javadi et al., 2017) and with detour demands (Javadi et al., 2019). It seems likely for taxi drivers the prefrontal cortex would be engaged when it is difficult to exploit a hierarchy, such as instances of planning routes circuitous routes within a region where no hierarchy can be exploited. Future work could adapt our methodology for brain imaging studies in order to clarify the neural structures involved in expert planning.

### Conclusion

The current study provided real-world evidence of hierarchical route planning. The vast complexity of existing street networks, such as London, renders these environments computationally intractable for traditional tree-search models of situated route planning. Hierarchical models, which segregate the environment and represent it through smaller clusters containing local information (McNamee, Wolpert & Lengyel, 2016), are a promising approach, but it has been challenging to find real-world evidence for them. Previous work had employed virtual environments, which have hierarchically organised clusters by design, in contrast to urban networks, which often lack clearly defined boundaries. Here, we bridged this gap by relying on agreement rates across taxi drivers as an indicator for boundaries. We were then able to test the impact of boundaries and of other street network properties on route recall tasks performed by London taxi drivers. Our ecologically valid data shows that boundaries impact response times during route recall, as do other street network properties such as turning actions and road types. Taken together, our findings support the view that taxi drivers employ a hierarchical representation of London when planning routes.

## Conflict of Interest Statement

The authors have no conflicts of interest to declare

## Acknowledgements.

We thank the Engineering and Physical Sciences Research Council UK (EPSRC) and Ordnance Survey Ltd for funding our research. The British Academy supported the work of PFV (PFSS23\230053). We thank Iva Brunec for her help preparing figures.

## Notes

### Competing Interest Statement

The authors have declared no competing interest.

