## Supplementary Materials for "London taxi drivers exploit neighbourhood boundaries for hierarchical route planning"

| Distance (km) | | | | | | |
|---|---|---|---|---|---|---|
| **Study** | **No** | **Start Location** | **Goal Location** | **Euclidean** | **Shortest Path** | **Cardinal Direction**[1] |
| 1 | 1 | Chelsea Harbour<br>*Chelsea Harbour Drive* | London Heliport<br>*Bridges Court* | 0.853 | 2.863 | 161 |
| 1 | 2 | The Radisson Edwardian Sussex Hotel<br>*Granville Place* | The Grange Fitzrovia Hotel<br>*Bolsover Street* | 1.269 | 1.813 | 47 |
| 1 | 3 | Putney Bridge Station<br>*Ranelagh Gardens* | Harrods<br>*Brompton Road* | 4.669 | 5.610 | 42 |
| 1 | 4 | The Kuwait Health Office<br>*Devonshire Street* | Selfridges<br>*Orchard Street* | 0.984 | 1.318 | 208 |
| 1 | 5 | The Tate Modern<br>*Holland Street* | Moorgate Station<br>*Moorgate* | 1.518 | 2.602 | 36 |
| 1 | 6 | Marlborough London Arts Gallery<br>*Albemarle Street* | Buck's Club<br>*Clifford Street* | 0.270 | 0.596 | 358 |
| 1 | 7 | Savoy Circus<br>*Westway* | Old Street Station<br>*Old Street* | 11.172 | 12.272 | 83 |
| 1 | 8 | Joe Allen's Restaurant<br>*Exeter Street* | The Doubletree Courthouse Hotel<br>*Great Marlborough Street* | 1.375 | 2.087 | 282 |
| 1 | 9 | Stockwell Station<br>*Clapham Road* | The Royal Hospital in Chelsea<br>*Royal Hospital Road* | 3.024 | 4.539 | 306 |
| 1 | 10 | The Savoy Hotel, River Entrance<br>*Savoy Place* | The Royal Society of Arts<br>*John Adam Street* | 0.242 | 1.409 | 260 |
| 1 | 11([2]) | Cutty Sark<br>*King William Walk* | Island Gardens Station<br>*Manchester Road* | 0.626 | 7.698 | 353 |
| 1 | 12 | The Home Office<br>*Marsham street* | Monument Station<br>*King William Street* | 3.367 | 4.159 | 61 |
| 2 | 8[3] | Bill's<br>*Exeter Street* | Double Tree Courthouse Hotel<br>*Great Marlborough Street* | 1.375 | 2.087 | 282 |
| 2 | 13 | Royal Oak<br>*Lord Hills Bridge* | Revolution<br>*Clapham High street* | 7.248 | 9.490 | 149 |
| 2 | 14 | Zizzi St Giles<br>*Bucknall street* | The Gate Theatre<br>*Pembridge Road* | 4.909 | 5.399 | 261 |
| 2 | 15 | Maudsley Hospital<br>*Denmark Hill* | Shoreditch Park<br>*Mintern Street* | 7.075 | 8.291 | 4 |
| 2 | 7 | Savoy Circus,<br>*Westway* | Old Street Station<br>*Old Street* | 11.172 | 12.272 | 83 |
| 2 | 16 | Elephant and Castle Stn<br>*Elephant Rd* | Swiss Cottage<br>*Finchley Road* | 7.377 | 8.959 | 316 |
| 2 | 17 | Tower of London<br>*Tower Hill* | Holland Park Car Park | 8.399 | 9.063 | 264 |
| 2 | 18 | Euston Station<br>*Exit on Eversholt Street* | Holborn Station<br>*Holborn* | 1.436 | 1.450 | 146 |

1. Direction of the goal location from origin; N = 0°; E = 90°; S = 180°; W = 270°
2. Removed from data analysis due to invalid recall format
3. Change of name

**Table S1.** Geographical Properties of Study Tasks.

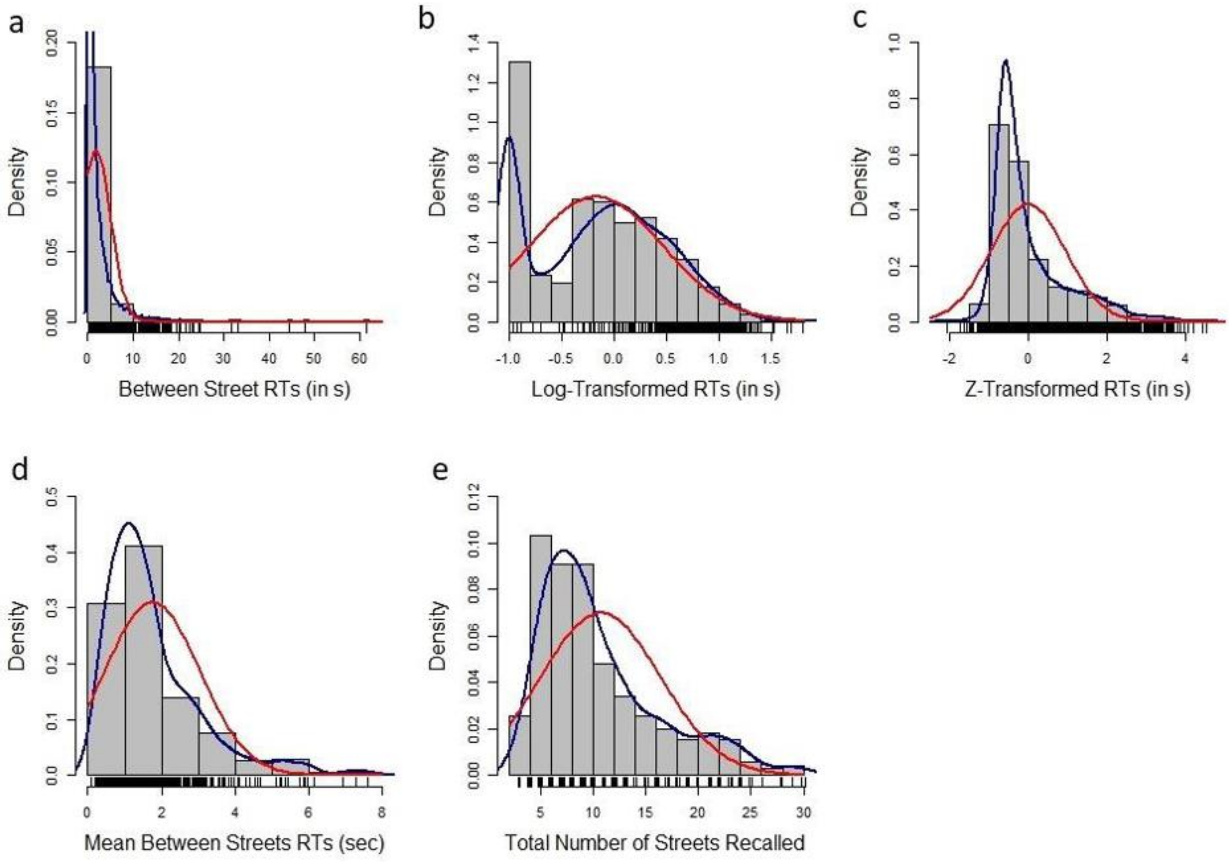


**Figure S1.** Summary of response time distributions. Distributions of initial response times for the raw (a), the log-transformed (b) and the z-standardized (c) data, as well as the mean of the response times between streets per route and driver (d) were plotted. Figure (e) visualises the total number of streets that were recalled. Fitted distribution (blue line) and normal distribution (red line) indicate skewness of the raw data (a) and a best fit for log-transformed RTs (b) over z-standardised RTs (c). The fitted distribution (blue line) of raw between street RTs was cut off as it exceeded the plot beyond y-scale limits. Note: The cut off point for transcription of fast response times between streets was at 0.1 sec (fastest RTs).

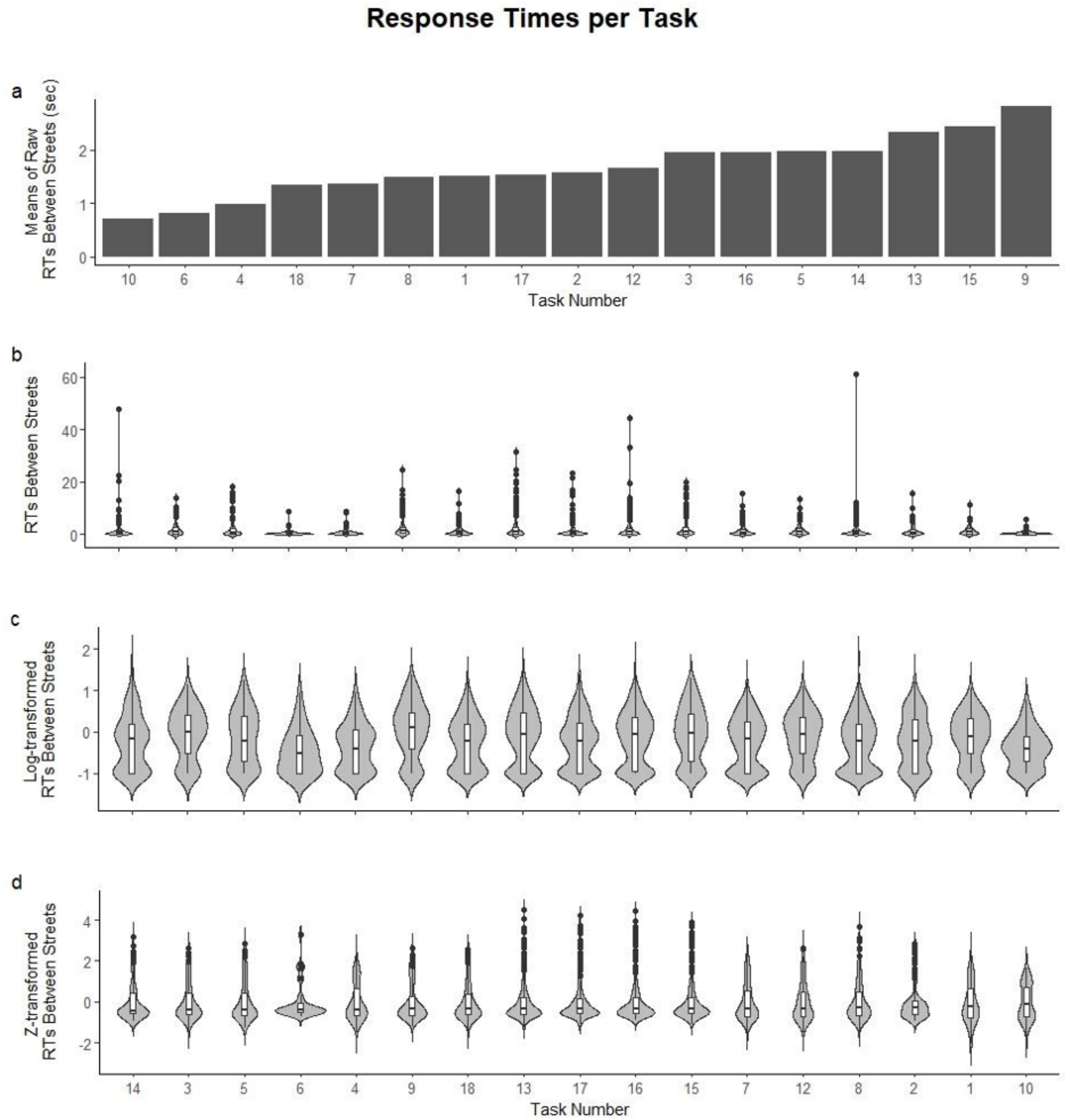


**Figure S2.** Response times for each route. (a) Bar plot of the mean response times between streets for each route by speed in ascending ordered (0.7s for route 10 to 2.8s for route 9). Violin plots of the raw (b) and z-transformed (d) RTs between streets indicate a range of response times outside the interquartile range for all runs. Violin plots of the log-transformed data (c) highlight the density distribution of data for each run. Cf. Figure 4.4

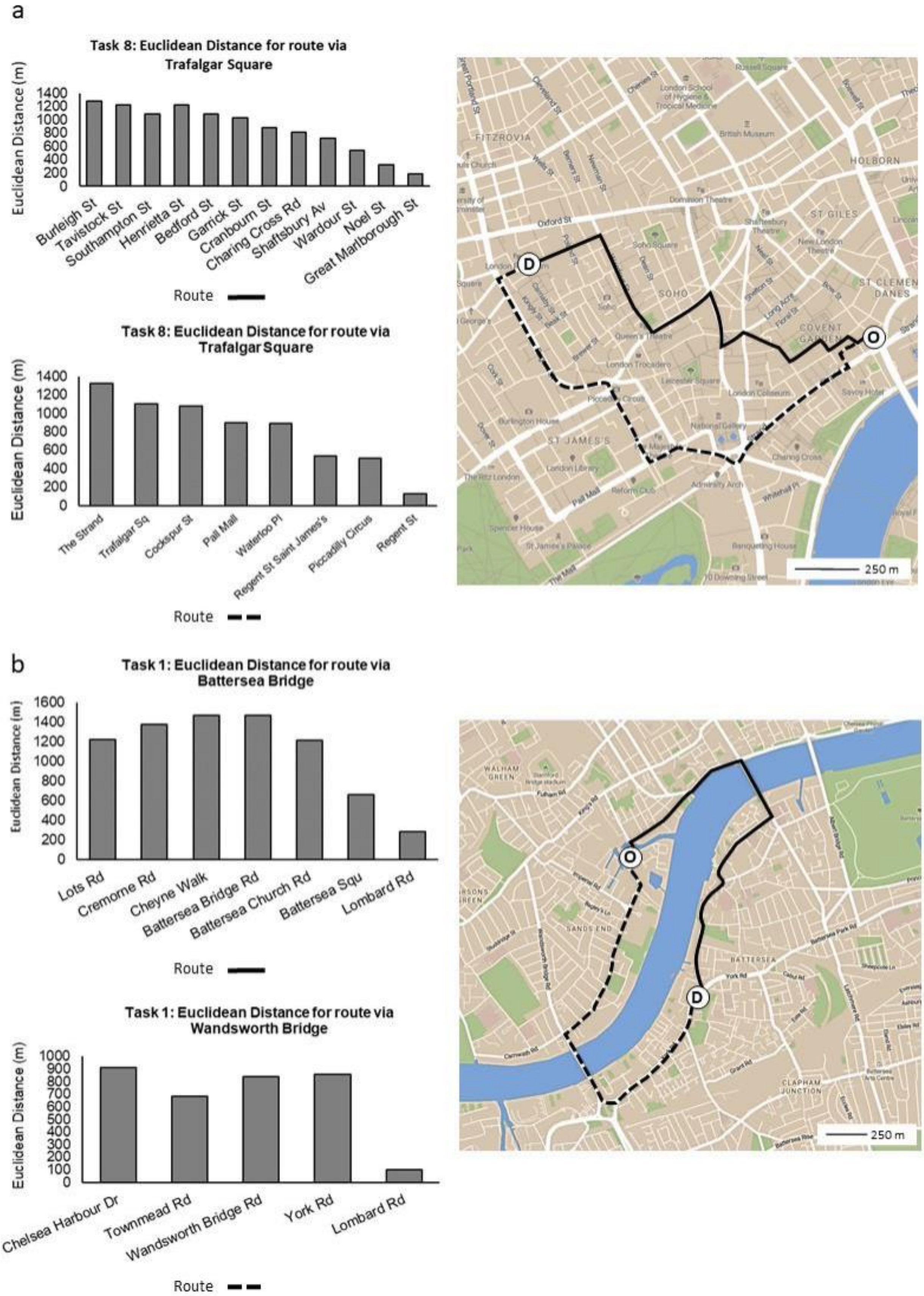


**Figure S3.** Circuity values. Euclidean distances measured (left) and mapped runs (right) show two different routes with no detour (a) and a forced detour (b). Euclidean distance along individual streets in routes that were approaching the destination directly constantly decrease (a), whereas routes with detour character (b) show stages of increasing Euclidean distance a decrease occurs closer to the destination.

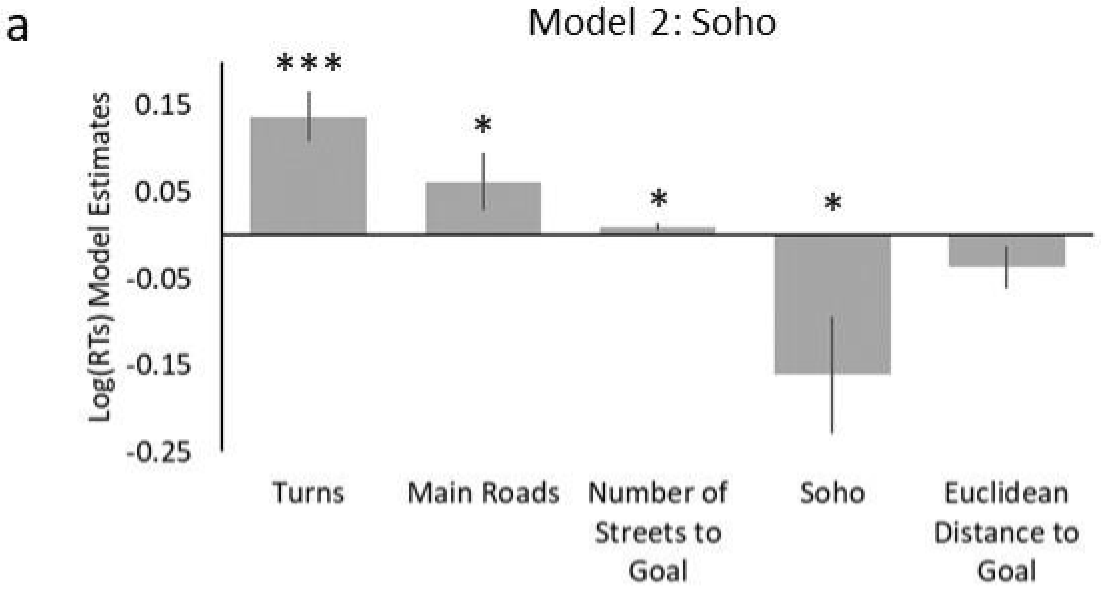


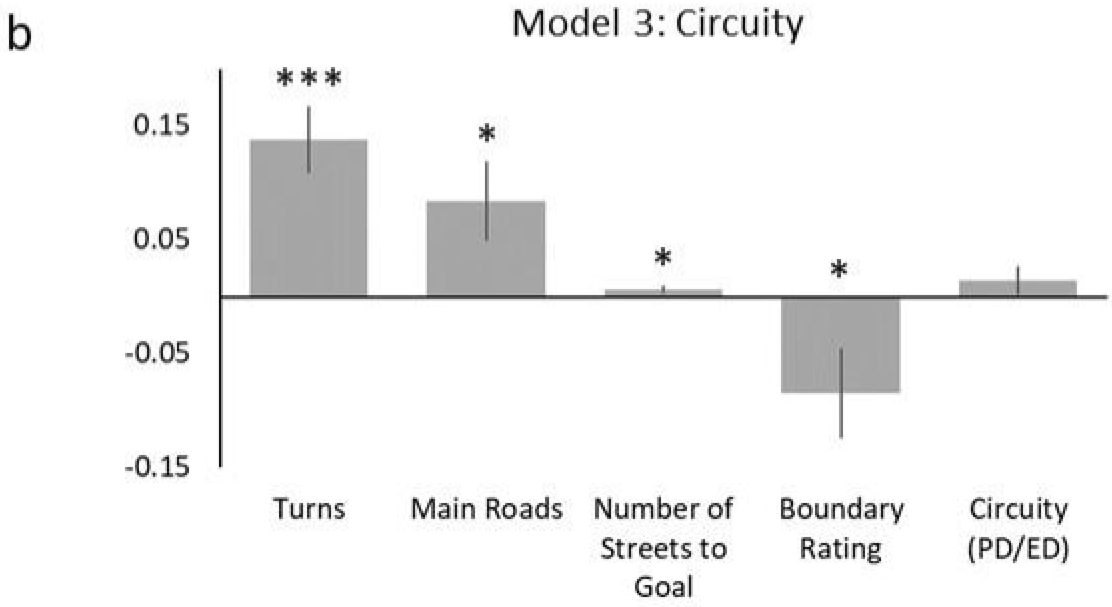


**Figure S4.** Parameter estimates for alternative models. Two alternative models were analysed in addition to the basic model. In the first alternative model (Model 2), boundary agreement rates were replaced by Soho including its boundaries. In Model 3, Euclidean distance was replaced by circuity, a distance measure with detour character. Both models are in line with the original model and tendencies remain the same.

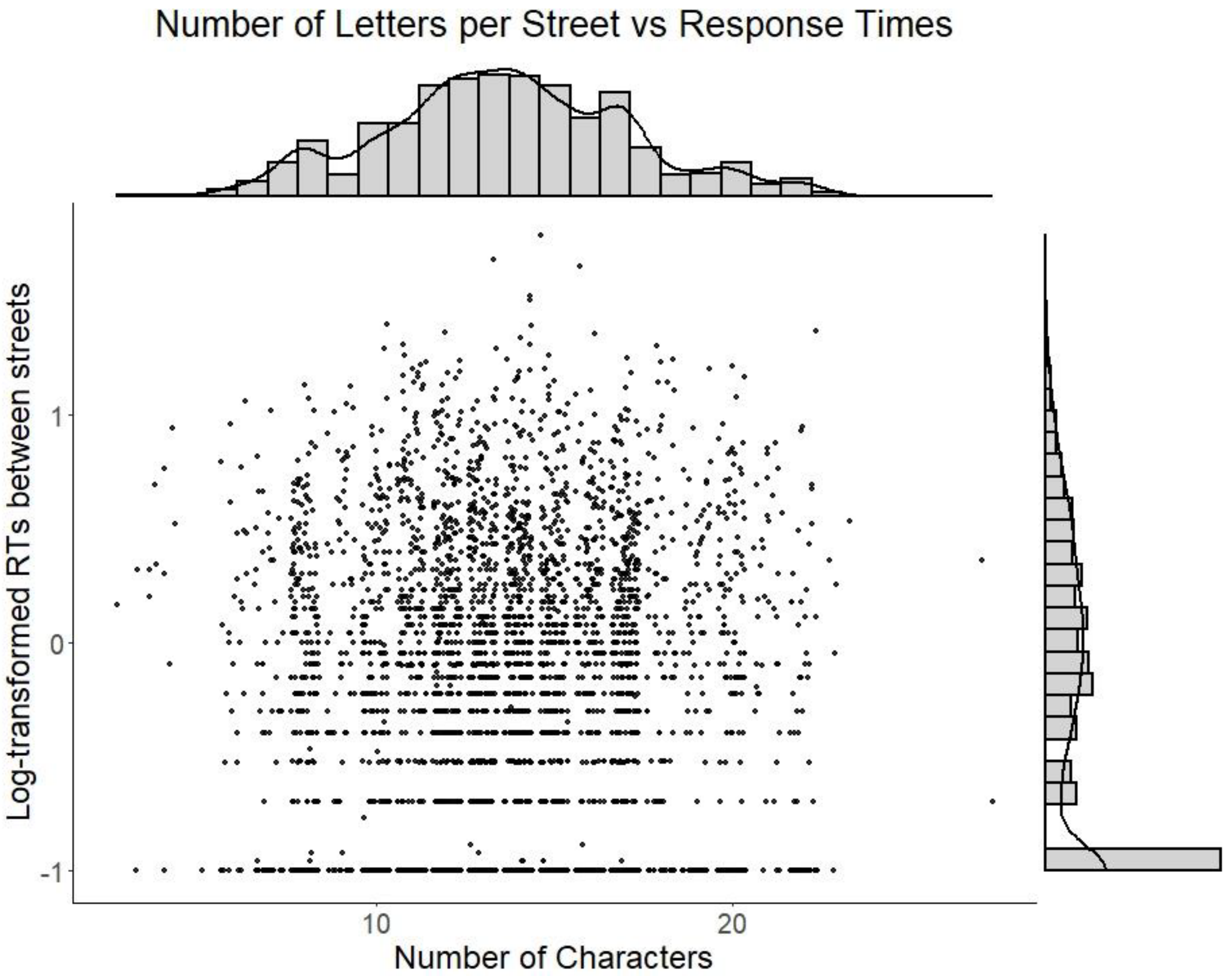


**Figure S5.** Relation between linguistic measures and response times. To account for a potential linguistic impact of the word length on the recall of individual streets, the number of letters were plotted in relation to log-transformed response times between streets. There was no relation between the two variables.
